# A closer look at plankton: potential interactions inferred from centimeter-scale in situ observations

**DOI:** 10.64898/2026.05.18.725820

**Authors:** Thelma Panaïotis, Jean-Olivier Irisson, Mara Freilich, BB Cael

**Affiliations:** National Oceanography Centre, SO143ZH Southampton, UK; Laboratoire d’Océanographie de Villefranche, Sorbonne Université, 06230 Villefranche-sur-Mer, France; Department of Earth, Environmental and Planetary Science, Division of Applied Mathematics, Brown University, 02912 Providence, RI, USA; Department of the Geophysical Sciences, University of Chicago, 60637 Chicago, IL, USA

**Keywords:** Plankton distribution, Spatial pattern, Microscale, In situ imaging, Ecological network

## Abstract

Plankton are essential to marine ecosystems, supporting food webs and mediating biogeochemical processes such as carbon export to depth. Their spatial distribution influences ecosystem dynamics and serves as an indicator of environmental change. Although drifting plankton could be expected to exhibit random distribution, numerous studies have revealed significant heterogeneity in their spatial patterns. However, very few studies targeted plankton distribution at the centimeter scale in situ, despite its importance for understanding biological processes. We argue that centimeter-scale distances in plankton could reveal potential ecological interactions. Using an extensive in situ dataset of 18 million planktonic organisms collected by the In Situ Ichthyoplankton Imaging System (ISIIS), which images multiple organisms simultaneously and preserves their positions in the water column, we analyzed centimeter-scale distances in plankton. By comparing observed distances with those expected under a random distribution, we assessed potential interactions at three levels: among all organisms, within plankton groups and across groups. Our results show that planktonic organisms exhibit non-random distributions at the centimeter scale, with smaller distances than expected, suggesting potential ecological interactions. Notably, distances up to 11 cm were the most informative, which is much larger than typical interaction distances in plankton. Additionally, observed distances were compatible with a simple attraction model. Finally, we propose the non-randomness of distances as a novel metric of interaction strength in plankton ecological networks and compare it against classical empirical or co-occurrence networks. These results offer new insights into in situ interactions and how they shape plankton distribution at centimeter scale.

**Significance statement:** This study reveals that planktonic organisms exhibit non-random spatial distributions at the centimeter scale, highlighting the importance of ecological interactions in shaping their distribution at this scale. By analyzing an extensive in situ plankton imaging dataset, we introduce a novel metric of interaction strength based on the non-randomness of distances between organisms, and compare it to common interaction metrics. These findings challenge the traditional view of plankton as passive drifters by highlighting that their distribution at microscale is shaped not just by physical processes such as turbulence but also by ecological interactions.

**Author contributions:** JOI contributed to data acquisition. TP processed the data under the supervision of JOI. TP, JOI and BBC designed the study. TP conducted the analyses under the supervision of MF, JOI and BBC. TP wrote the initial draft of the manuscript. All authors contributed to the interpretation of results, supported manuscript preparation and approved the final submitted version.

## Introduction

Interactions between organisms are fundamental drivers of ecosystem dynamics, playing a key role in the maintenance and evolution of life on Earth. Ecological networks provide a powerful framework for representing and analyzing these interactions Bascompte (2009). However, detecting interactions usually requires considerable sampling efforts Kéfi et al. (2015). As a consequence, interactions are generally inferred from indirect evidence, including phylogenetic relationships, function traits or spatial distribution Morales-Castilla et al. (2015). This has led to the development of various methods to infer interactions from patterns in observational data.

Among these methods, co-occurrence networks Proulx et al. (2005) are used to infer interactions based on statistical correlations, assuming that positive correlations indicate interactions such as predation or mutualism, while negative correlations may suggest avoidance. However, this approach tends to confound true ecological interactions with shared environmental preference Freilich et al. (2018). Notably, recent work has shown that complex co-occurrence patterns can arise from very sparse underlying interaction networks Camacho-Mateu et al. (2024). Furthermore, when the spatial positions of individuals are known, spatial point pattern analysis (SPPA) becomes a powerful tool for inferring potential ecological interactions Velázquez et al. (2016); Ben-Said (2021). Indeed, various ecological processes (e.g. biotic interactions, dispersion) can influence the spatial distribution of species, and these patterns can be detected using SPPA Legendre and Fortin (1989). Originally developed in forestry, SPPA has been mostly used in plant ecology Velázquez et al. (2016) but has also been applied to animal-related features Klaas et al. (2000). While SPPA is well suited for studying stationary organisms or features, its application becomes more challenging in dynamic environments, such as marine ecosystems.

Planktonic organisms – defined as drifters unable to swim against currents – span a wide range of taxonomic groups and sizes, from micrometers to meters. Photosynthetic phytoplankton are primary producers, while zooplankton are a trophic link between phytoplankton and higher trophic levels. Plankton also contribute to the export of organic carbon to deeper waters via the biological carbon pump Falkowski (2012). Despite being drifters, plankton distribution is heterogeneous across various spatial scales Davis et al. (1992); Benoit-Bird et al. (2013); Robinson et al. (2021). This patchiness can result from physical forces like density gradients (e.g. fronts, stratification) Prairie et al. (2012) or biological processes such as blooms Behrenfeld and Boss (2014). These patterns typically occur at scales of tens of meters to hundreds of kilometers, orders of magnitude larger than the scales at which biological interactions take place. At these scales – millimeter to centimeter – heterogeneity can arise from behavioral activity such as swimming Gallager et al. (2004), turbulence, or their interplay Prairie et al. (2012). Indeed, many zooplanktonic organisms swim actively to encounter food and mates, making motility a critical aspect of their fitness Visser (2007). Swimming speed depends on predation risk and food availability, but also scales with organism size Kiørboe (2011). The study of these fine-scale behaviors require tools that can resolve spatial distributions at the scale of the organisms.

The advent of in situ imaging makes it possible to observe plankton spatial distribution at the microscale Widder and Johnsen (2000); Gallager et al. (2004). Among other instruments, the In Situ Ichthyoplankton Imaging System (ISIIS) Cowen and Guigand (2008) offers the highest sampling rate (> 100 L s^-1^) and minimal disturbance to imaged organisms, allowing for the simultaneous detection of multiple organisms within the field of view. This allows the application of SPPA to in situ planktonic organisms, to understand the effect of ecological interactions on plankton distribution. Because fine-scale distance anomalies dissipates more rapidly with increasing distance than abundance anomalies, spatial proximity at organismal scales should offer a more sensitive indicator of ecological interactions than co-occurrence-based metrics.

In this work, we hypothesize that the physical distances between planktonic organisms at their interaction scale can reveal ecological interactions, because these interactions, such as predation, require physical proximity. To test this, we analyzed a large in situ imaging dataset. We measured physical distances between organisms at three levels – among all organisms, within plankton groups and across plankton groups (groups are listed in Table S1) – and compared them to distances expected under a random distribution. To further interpret the observed distances between planktonic organisms, we developed a simple attraction model to assess whether it could reproduce observations. Finally, the association metric derived from distances between organisms was compared against two other metrics, co-occurrence and size-based associations, to determine whether they capture similar ecological information.

## Results

### Imaged organisms were undisturbed

To assess whether turbulence generated by ISIIS affected planktonic organisms, we examined the posture and positioning of large (> 1 cm) and delicate organisms imaged by the imaging system. Their natural positions and intact structures (e.g. extended tentacles, no deformation) suggest that they were undisturbed (Figure S1). For smaller organisms, the image resolution and detail were insufficient to apply the same visual assessment. These observations suggest that disturbance from the imaging system is minimal and that centimeter-scale non-random distances between individuals are more likely shaped by behavior.

### Organisms are closer than expected under a random distribution

We focused on distances smaller than 11 cm, which was found to be the most informative range (Figure S2a). Our analysis reveals that distances between planktonic organisms – when considered all together, regardless of taxonomy – were different from those expected under a random distribution of planktonic organisms. In other words, the spatial distribution of planktonic organisms was non-random (Figure S3b). More specifically, planktonic organisms were closer to each other than randomly distributed particles would be (Figure 1a). This pattern was also true within 11 of the 14 plankton groups for which more than 10,000 distances could be measured (Figure S3c) and particularly pronounced for Acantharea, a subgroup of Rhizaria (Figure 1a). When considering pairs of plankton groups, distances were found to be non-random for about 3/4 of pairs for which enough distances were recorded (Figure S3d). Organisms were closer than expected except for one pair (Figure 1b), which consisted of Ctenophora and *Oithona* copepods. In our dataset, ctenophores were mostly carnivorous Cydippida, making this observation compatible with predation avoidance by *Oithona*. Overall, the strength of the non-randomness tended to increase with the number of available distances: when the sample size was sufficiently large, observed distances consistently deviated from those expected under a random distribution (Figure S3b-d). However, non-randomness was independent of the size of organisms (Figure S4).

**Figure 1:**
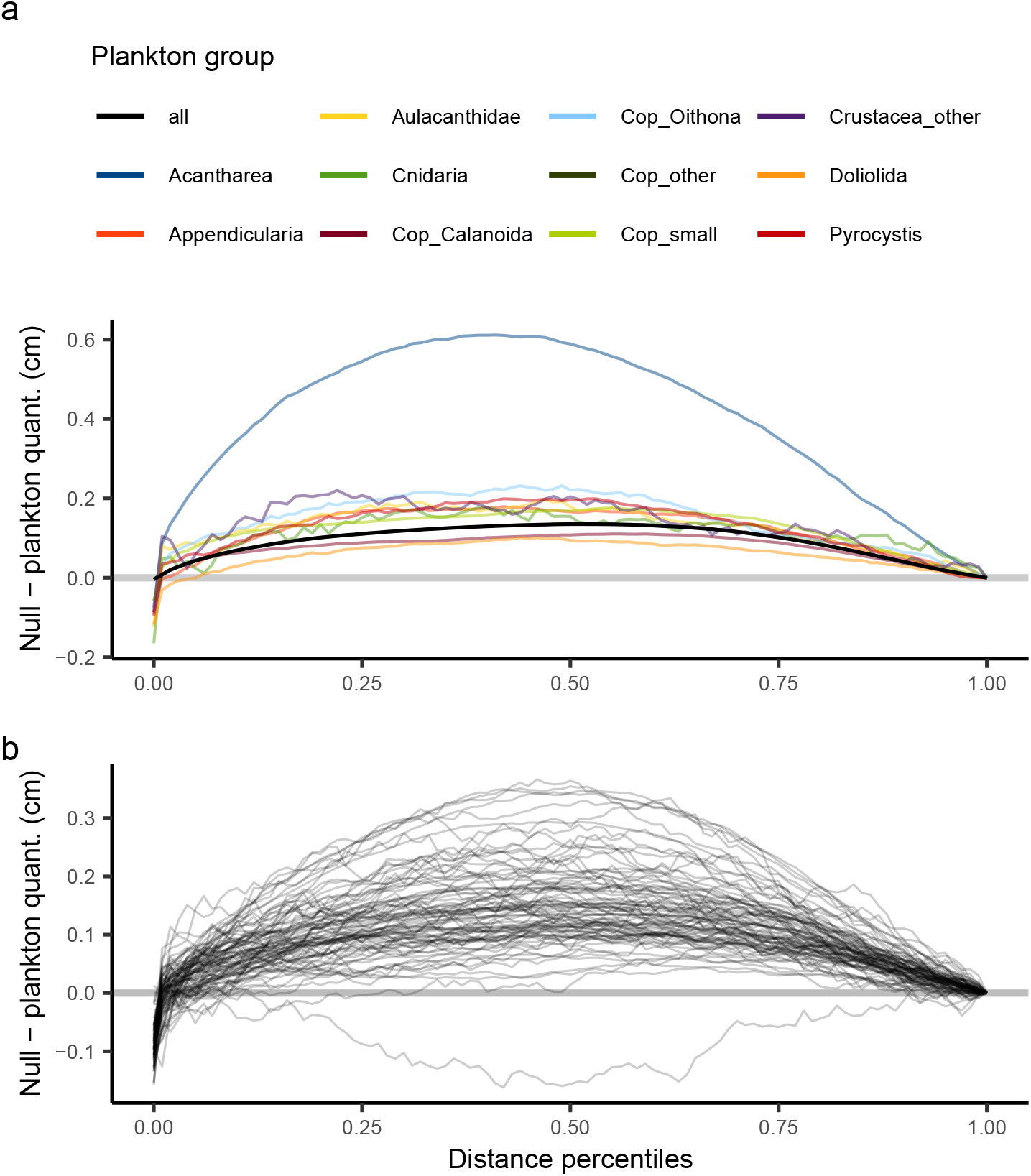
Difference between quantiles of expected distances (random distribution) and quantiles of observed plankton distances, plotted against distance percentiles. Positive *y* values indicate that organisms are closer together than would be expected by a random distribution, while *x* values indicate whether this affects smaller (left) or larger (right) distances. (a) shows all and intra-group distances, with colors representing plankton groups, (b) shows inter-group distances. Only cases where distances are significantly different from random are displayed.

### A simple individual-based model reproduced observed distances

To assess whether simple attraction between organisms could explain the observed distances, we developed a minimal individual-based model in which points were randomly distributed and then displaced once toward areas of higher density, simulating attraction. The model relied on a single free parameter – the strength of attraction – determined by a sensory radius and swimming displacement. Optimal values for these parameters were obtained via grid search: selected sensory radius was 10 cm and selected displacement length was 0.2 cm (Figure S5). Despite its simplicity, the resulting distance distribution closely matched the one observed in plankton (Figure 2). The model underestimated short distances, overestimated longer ones, and slightly underestimated the longest ones (grey dashed line), suggesting that real organisms may exhibit short-range repulsion not captured in the model. Nevertheless, this highly simplified approach reproduced key features of the distribution of observed distances.

**Figure 2:**
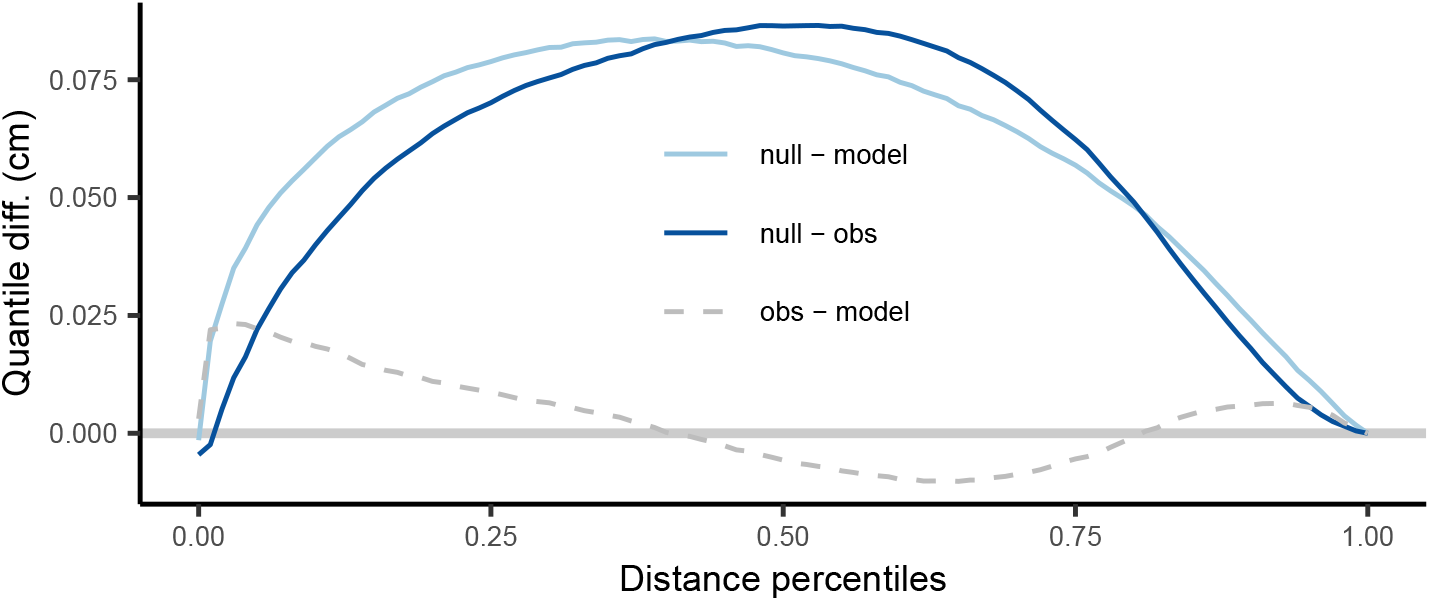
Difference between quantiles of expected distances (under random distribution), observed plankton distances and modeled distances obtained by the individual-based model, plotted against distance percentiles.

### Comparison with other metrics

Distances between planktonic organisms were shorter than expected under a random distribution, and this deviation could be explained by a simple attraction model. This suggests that in situ imaging is able to detect interactions between organisms. Building on this idea, we introduce a new distance-based association metric and compare it to two existing approaches. The first comparison metric is usually computed on empirical ecological networks, built from known relationships (e.g. predation); however, the limited taxonomic resolution of our dataset did not allow us to assign those relationships. Instead, we computed predation probability from size differences between plankton groups: this metric infers potential predator-prey interactions based on organism size. The second comparison metric, based on co-occurrence, is widely used to study plankton associations but has known limitations, such as the tendency to confuse shared environmental preferences with true ecological interactions Freilich et al. (2018). In contrast, our distance-based approach captures spatial proximity, which may more directly reflect behavioral processes. This comparison indicated that these three metrics do not necessarily capture the same association between plankton groups (Figure 3). In addition to Spearman correlations, the Mantel test revealed that association matrices (Figure S6) were indeed different (Table S2). Furthermore, when treating the size-based metric as an a priori hypothesis in a binary classification framework (i.e. presence or absence of association), the co-occurrence metric performed better at predicting expected associations than the distance-based metric (Table 1). More specifically, the former retrieved nearly all of the associations expected according to size (recall = 92.1%). Overall, these results highlight that these three metrics capture different aspects of the associations between plankton groups.

**Table 1:** Comparison of distance-based and co-occurrence metrics to the a priori hypothesis relying on the size-based metric. Accuracy corresponds to the proportion of correctly identified associations (both present and absent) out of all evaluated cases, precision corresponds to the proportion of identified associations that were actually expected, recall corresponds to the proportion of expected associations that were successfully identified.

| Metric | Accuracy | Precision | Recall |
| --- | --- | --- | --- |
| Distance | 35.3% | 59.5% | 18.3% |
| Co-occurrence | 67.0% | 70.0% | 92.1% |

**Figure 3:**
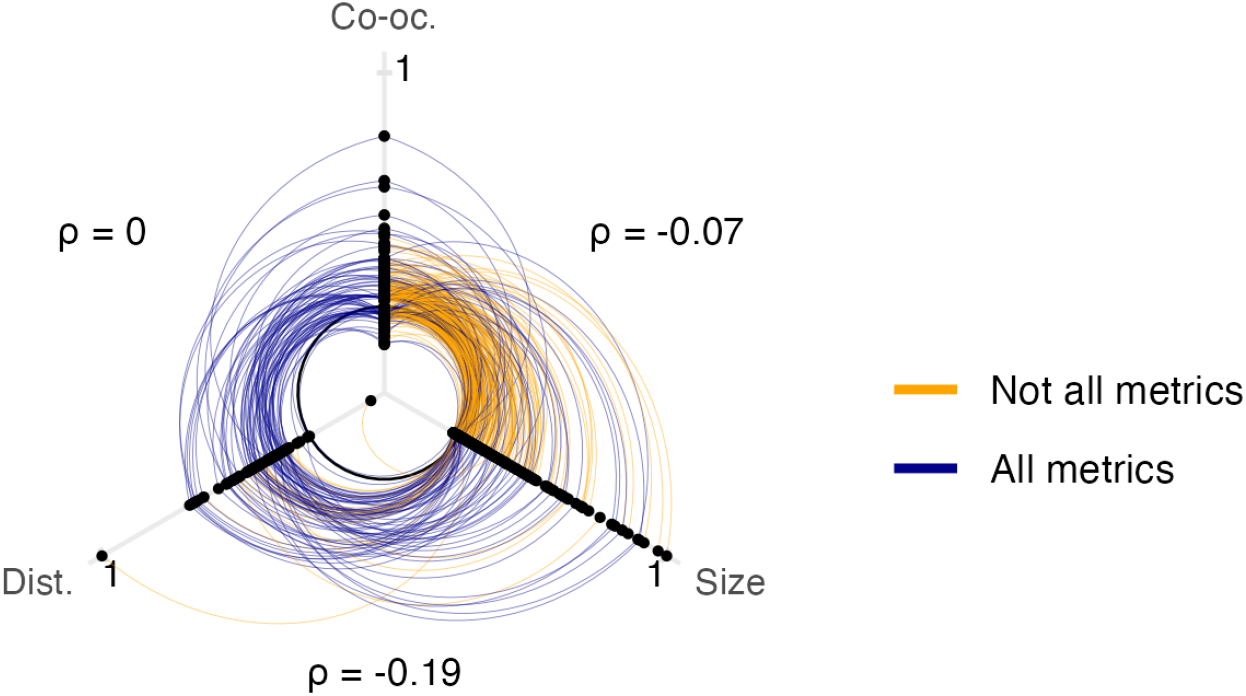
Comparison of association metrics. Black points represent association values for each pair and metric type, with lines connecting pairs across metric types. The black circle indicates the zero value for each metric. Some metrics, particularly the distance-based one, could not be computed for all pairs; this is reflected in the line color: blue lines connect points (i.e. group or pair of group) for which all 3 metrics could be computed; while orange lines connect points for which one metric is missing. Spearman correlation values (*ρ*) between metric types are also shown, all correlation p-values were > 0.05. Dist = distance-based, co-oc = co-occurrence. *ρ* = Spearman’s correlation coefficient.

## Discussion

### Summary of key findings on inter-individual distances

Overall, our results show that planktonic organisms tend to be more spatially clustered at the centimeter scale than would be expected under a random distribution. This pattern was true for inter-organisms distances regardless of plankton groups, but also for 80% of considered groups when focusing on intra-group distances; and also for 75% of group pairs when considering inter-group distances. This pattern depended on the number of available distances for each considered group or pair but not on the size of the organisms.

### Drivers of non-random spatial patterns in plankton

Non-random spatial patterns in plankton can result from various processes operating across multiple scales. At the mesoscale, physical structures such as fronts and eddies can create biological hotspots and lead to heterogeneous distributions, such as patches and filaments Owen (1981). At finer scales, the availability of resource patches can lead to local increases in density, but without necessarily producing non random distribution within patches. Similarly, shear associated with small-scale turbulence may create thin layers with elevated concentrations of organisms Durham and Stocker (2012); Prairie et al. (2012). At the microscale, turbulence may also cause aggregation and induce non-random spatial patterns among passive particles Gustavsson and Mehlig (2016); Pumir and Wilkinson (2016).

However, several lines of evidence suggest that turbulence is not the primary cause of the observed patterns. First, the images of undisturbed planktonic organisms imaged by ISIIS (Figure S1) suggest low turbulence in the instrument’s field of view. Second, if turbulence was the only driver of fine-scale distributions, one would expect to observe similar distances for groups (or pairs of groups) of similar sizes. This was not the case for, e.g., Acantharea and Appendicularia (median ESD = 0.88 and 0.93 mm, respectively). Regarding the potential effect of plankton thin layers, these are meter-scale features Prairie et al. (2012), larger than the scale of ISIIS images (∼10 cm in height, ∼50 cm in length). Thus, in an image captured within a thin layer, plankton concentration can be higher but can be considered isotropic. As the number of organisms in an image does not affect the distribution of distances between them (this affects only the distance to the nearest neighbor), this rules out the effect of plankton thin layers. In contrast, behaviors requiring close physical proximity are more likely to generate non-random inter-individual distances, particularly among motile organisms. Taken together, this supports a key role of behavior in shaping non randomness in plankton distribution at microscale.

Among plankton groups, Acantharea displayed much shorter inter-individual distances than expected under a random distribution. These mixotrophic Rhizaria are unicellular eukaryotes with strontium sulfate skeletons. In our observations, they were mostly found in surface waters (sometimes in patches spanning several meters of depth, Figure S7a), consistent with previous studies in the Ligurian Sea Faillettaz et al. (2016) and globally Decelle and Not (2015). However, as explained above, thin layers and microscale turbulence alone cannot explain these closer-than-random distances. To rule out classification errors (e.g. dead particles mistaken for Acantharea), we checked a subsample of 400 objects identified as Acantharea detected in patches and found that > 99% were correctly classified (Figure S7b), confirming that observed distances are not artefactual. These results thus indicate that behavioral mechanisms are likely responsible for the observed patterns. While non-motile, Acantharea can actively regulate their buoyancy Febvre-Chevalier and Febvre (1994), which could induce aggregation at specific depths. Acantharea sometimes feed on conspecifics Swanberg and Caron (1991), but our size-based metric predicted a low interaction probability (Figure S6). Another plausible behavioral explanation is reproduction: although Acantharea remain uncultivated and their reproduction is poorly understood, they release “swarmers” thought to be gametes Decelle and Not (2015), and spatial proximity could facilitate fertilization.

Furthermore, the greater distances observed between Ctenophora and *Oithona* are consistent with the avoidance behavior of copepods toward carnivorous Cydippida, who capture prey using two long tentacles Nival et al. (2020). While the co-occurrence analysis did not detect a significant spatial correlation between Ctenophora and *Oithona*, our size-based metric supports the potential predation of *Oithona* by Ctenophora (Figure S6). In our dataset, other groups of copepods were present (some bigger, some smaller), but *Oithona* were the only one showing this pattern. This difference could be due to different swimming or sensory ability. The reported escape distances in copepods are generally < 10 mm Buskey et al. (2002), much smaller than our observed distances. This suggests that this mechanism is not sufficient to explain observed distances. Instead, *Oithona* might be able to detect ctenophores from a greater distance – similar to how copepods can sense marine snow trails over several centimeters Lombard et al. (2013) – and adapt their swimming to avoid encounters with ctenophores. Finally, our segmentation algorithm often missed the tentacles of ctenophores and detected only the body, meaning distances are typically computed between the copepod and the ctenophore body centroid. In such cases, a copepod could be much closer to the tentacles than the measured distance suggests.

### Comparison with previous work and theory

These results contrast with those reported by Widder and Johnsen (2000), who investigated the 3D distribution of dinoflagellates and copepods in situ. By focusing on nearest-neighbor distances, they found no significant deviation from randomness for dinoflagellates at low abundances, but a statistically significant difference emerged at higher abundances with organisms being further away than expected under random distribution. For copepods, their observations were compatible with a random distribution. At the meter scale, patchiness was found to vary considerably across plankton groups, with smaller organisms forming larger and looser patches Robinson et al. (2021). Based on these findings, one might expect smaller organisms to exhibit larger inter-individual distances. Surprisingly, our analysis of intra-group distances revealed no clear relationship between organism size and either inter-individual distances or the non-randomness of these distances (Figure S4), suggesting that the spatial scale of interactions does not systematically vary with organism size within the considered size range.

In terms of ecological interpretation, our findings diverge from the expectations of the ideal free distribution theory (IFD), which posits that organisms within a taxonomic group distribute themselves in space to minimize competition for resources Fretwell and Lucas (1969). At large spatial scales, this can lead to uneven density distribution, e.g. herbivores concentration around patches of phytoplankton. However, at smaller scales, resource distribution tends to come more homogeneous, especially at the centimeter scale we investigate in this work. Under these conditions, one would expect a relatively uniform spatial distribution, with greater distances between individuals than would occur under random distribution. Yet, the intra-group distances we observed were not only shorter than those under a uniform distribution, but also shorter than expected under a random distribution. This suggests that the assumptions of the IFD may no longer hold at this scale.

Regarding inter-group distances, predator-prey interactions could contribute to shorter observed distances. Indeed, a predator would be expected to swim towards a prey, while the prey would swim away from the predator. Since predators are generally larger and swim faster than their prey Hansen et al. (1994); Kiørboe (2011), this could result in shorter distances between the two groups. However, not all groups in our dataset represent clear predator-prey pairs, and one counterexample (Ctenophora and *Oithona*) displays distances larger than expected despite being potential predator and prey. Furthermore, due to the limited taxonomic resolution, a given group may include a mix of different trophic strategies. Thus, while predator-prey interactions offer a plausible explanation for our observations, they are unlikely to account for all cases.

Our analysis identified an optimal distance threshold of 11 cm which maximized the difference between observed plankton distances and random distances (Figure S2a). Despite this value being close to the ISIIS image height (10.5 cm), this is unlikely to be related, as ISIIS images have a small pitch due to the oscillating tow pattern and plankton distribution at centimeter scale is likely isotropic. Furthermore, although this optimal threshold was identified using all organisms regardless of taxonomy, it also held for most cases of intra and inter-group distances (Figures S2bc), suggesting that the distance of potential interaction does not necessarily scale with organism size. Beyond this threshold of 11 cm, observed distances increasingly resembled random ones, suggesting a randomization of spatial patterns at larger scales. This implies that interactions between planktonic organisms likely diminish (or are absent) at distances greater than ∼10 cm. Interestingly, this threshold is substantially higher than typical interaction distances reported for plankton. For example, interaction ranges are estimated at 0.7-1.2 cm for fish larvae Hunt von Herbing and Gallager (2000), 3 mm for chaetognaths Feigenbaum and Reeve (1977) and 0.1-0.7 mm for *Acartia* copepods Jonsson and Tiselius (1990). Indeed, detection ranges are generally expected to be on the order of the body size Kiørboe (2011). However, previous work suggests that plankton may perceive each other at much greater distances than traditional behavioral experiments indicate: Haury and Yamazaki (1995) reported that the distance to the nearest neighbor in copepod aggregations often far exceeded known perception ranges; Doall et al. (1998) observed that male *Temora* copepods were able to detect females at distances of up to 13 cm. Our findings align with these observations while coming from direct in situ observations, and are further supported by our individual-based model, which successfully reproduced the observed spatial distributions using a detection range of 10 cm.

### Attraction-based modeling of plankton distances

Our individual-based model, which simulates a simple attraction behavior, closely reproduced observed plankton distances using only two parameters: a sensory radius of 10 cm (which aligns well with the identified threshold of 11 cm) and displacement length of 0.2 cm. However, the simplicity of the model introduces some limitations: it does not account for repulsion at very close range, likely explaining its slight underestimation of shorter distances, while more realistic individual-based models of collective behavior typically rely on both attraction and repulsion rules (e.g. Couzin et al. (2005)). Furthermore, our model simulates only a single iteration, without accounting for temporal evolution. It also neglects the influence of resource distribution, which is essential in shaping plankton distribution. Despite these limitations, our findings support the idea that simple behavioral rules can give rise to complex spatial patterns, consistent with previous studies DeAngelis and Mooij (2005); Berdahl et al. (2013). This suggests that microscale plankton distribution can emerge from basic interaction rules.

### Comparison between association metrics

Finally, our results highlight that the different association metrics assessed here (distance-based, size-based and co-occurrence) capture different ecological information. When using the size-based metric as an a priori hypothesis for potential predation interactions, the co-occurrence metric more effectively retrieved the expected associations than the distance-based metric. Nonetheless, co-occurrence networks are influenced by both environmental conditions and species interactions, without a clear means of disentangling these effects Freilich et al. (2018). In contrast, we argue that snapshot spatial distances may offer a more direct and dynamic signal of interaction potential, especially for processes like predation or mutualism that require physical proximity. By capturing the immediate spatial relationships among organisms, the distance-based metric may better reflect the mechanisms underlying ecological interactions, rather than their statistical correlations.

In conclusion, we demonstrate that the distances between planktonic organisms at the centimeter scale are significantly smaller than expected under a random distribution, indicating potential ecological interactions. We introduce this non-random spatial distribution of organisms as a novel metric for assessing association strength, offering complementary insights to those provided by traditional methods. Furthermore, these distances could be explained by simple attraction among individuals. These findings provide new perspectives on in situ interactions at the microscale by challenging the notion of plankton as passive drifters: at centimeter scale, plankton distribution is not only shaped by physical processes such as turbulence but also by ecological interactions enabled by their swimming abilities. Our results also suggest that planktonic organisms may possess the ability to detect each other over greater distances than previously thought.

## Material and Methods

### Plankton Imaging Data

Data was collected in the Ligurian Sea (NW Mediterranean) during the VISUFRONT campaign in the summer of 2013. Plankton imaging data was collected using the In Situ Ichthyoplankton Imaging System (ISIIS) Cowen and Guigand (2008), a high sampling rate (> 100 L s^-1^) in situ imaging instrument towed behind a ship and that oscillates between the surface and a given depth (here ∼100 meters). ISIIS used shadowgraphy to achieve a large telecentric field of view (10.5 cm length × 10.5 cm height × 50 cm depth, i.e. ∼5.5 L imaged per *frame*), minimizing parallax errors. Because imaging relies on a line-scan camera, collected frames can be assembled to generate a continuous ribbon of frames, which can be arbitrarily segmented to produce individual *images*. These images were set to be approximately 50 cm long to maximize the simultaneous detection of multiple planktonic organisms in a single image. Due to the oscillating sampling pattern, images are not perfectly horizontal but have a pitch of a few degrees, which reduces directional bias in measured distances. Finally, the large field of view of ISIIS relative to the size of the imaged organisms allows plankton to be observed with minimal disturbance, as evidenced by the natural positioning of delicate gelatinous organisms (Figure S1).

Given the large volume of data collected (∼80h of recording at > 100 L s^-1^), the processing pipeline was fully automated. After flat-fielding of images, planktonic organisms were detected using an intelligent segmentation method based on Convolutional Neural Networks (CNN), with a detection rate of 92% Panaïotis et al. (2022). Segmented objects were then sorted into taxonomic and/or morphological groups using a CNN classifier Panaïotis et al. (2025). A probability threshold was applied to classification scores to aim for a 90% precision rate, at the cost of a lower recall Faillettaz et al. (2016). In the end, 27 groups met the targeted precision rate (assessed on an independent test set) and were retained, with classification precision and recall presented in Table S1.

The resulting dataset contained ∼18 million planktonic organisms across ∼600,000 images with at least one detected organism; with a number of organisms per image ranging from 1 to 547. For each organism, the dataset includes its plankton group, position within the image (based on its centroid) and size computed as equivalent spherical diameter (ESD).

### Association metrics

From this dataset, three association metrics were computed: distance-based, co-occurrence and size-based.

#### Distance-based association

We computed physical distances between planktonic organisms detected in ISIIS images, using the centroid positions of the organisms. Distances were analyzed at three levels: between all organisms (irrespective of taxonomy), within plankton groups (intra-group) and across plankton groups (inter-group). This resulted in 3×10^8^ distances when considering all organisms, but the number could drop below 100 for poorly represented classes (e.g. *Pleuromamma* copepods). To ensure statistical robustness, we only considered plankton groups or pairs with at least 10,000 computable distances. For computation efficiency in cases with more than 10,000 distances, we summarised distributions using 10,000 quantiles. However, with ISIIS, the horizontal pixel size can be different from the vertical one depending on the scanning rate and towing speed (e.g. if scanning rate is too high compared to towing speed, the effective horizontal pixel size will be shorter than the vertical one and organisms will appear stretched vertically). This artifact was quantified by using spherical organisms (solitary Collodaria, belonging to Rhizaria, Figure S8): their deformation from a perfect circle was used to globally correct the horizontal positions of organisms in all images. Because ISIIS provides organism position in only two dimensions, with no information on the third spatial axis (i.e. the depth of the field of view, 50 cm), we tested whether 2D distances were representative of true 3D distances. To do this, we conducted a simulation experiment where both 3D and 2D distances were computed for a set of 1000 images where points were randomly drawn in a 3D space representative of the volume imaged by ISIIS. 2D and 3D distances were indeed highly correlated (R^2^ = 85%), supporting the use of 2D distances as a reasonable proxy for spatial proximity.

To assess whether distances between planktonic organisms carry ecological information, they were compared to those expected under a random distribution of organisms. To compute these random distances, a null dataset was created with the same properties as the plankton dataset, but with organism positions randomly generated within 2D image boundaries. This preserved both imaging area constraint and organisms count but removed any biological structure. Comparison between the distributions of plankton and null distances was performed with the Kuiper statistic (plankton to null Kuiper statistic, referred to as PNKS hereafter) on 10,000 quantiles of computed distances.

To focus on ecologically relevant interactions, we excluded distances that were too large to reflect ecological interactions, which occur at the scale of the organisms. We identified an upper distance threshold using a signal-to-noise ratio (SNR) approach: PNKS was computed for a range of distance thresholds (with null distances constrained by the same threshold) and the threshold yielding the highest PNKS was selected as the most informative. The detected threshold when considering all planktonic organisms was 11 cm (Figure S2a). To assess whether this threshold was group-dependent, the same approach was conducted within plankton groups and a majority of them returned the same optimal threshold of 11 cm (Figures S2bc). Thus, only distances shorter than 11 cm were retained; and only cases with at least 10,000 such distances were included in the analysis.

The computed PNKS alone was not informative because it depends on factors such as the number of computed distances. To determine whether a given PNKS value could indicate an association, we compared it to the Kuiper statistic obtained from comparing two null datasets (i.e. datasets where the positions of organisms are drawn randomly) – referred to as the null to null Kuiper statistic, NNKS – using 10,000 quantiles. This process was repeated pairwise with 20 null datasets, resulting in 190 NNKS to obtain a distribution of expected KS values for null dataset comparisons. Additionally, since NNKS itself depends on the number of distances, this computation was repeated across subsampled null datasets of various sizes (Figure S3a). Due to the computational cost of distance calculation, the maximal number of distances to compute in this experiment was set to 10^7^. To estimate NNKS for larger datasets, we extrapolated the relationship between NNKS and number of distances using quantile regression (5% and 95%) after log-transformation of both axes. To verify this extrapolation, we computed the NNKS values for 3 datasets with 10^8^ distances and confirmed consistency with extrapolated values (Figure S3a). We also tested whether using a global null dataset representative of all planktonic organisms regardless of taxonomy was appropriate for specific sets of distances (e.g. intra-group distances). For this, 5 null datasets restricted to Acantharea only were generated using the same method as above. Obtained NNKS values were similar to those obtained when considering all distances, validating the use of global null datasets for all analyses (Figure S3a).

Significant differences in PNKS values compared to the NNKS distribution indicated a non-random distribution of the planktonic organisms considered (Figure S3b-d). In such cases, the 10,000 quantiles of both plankton and null distances were extracted and used to compute differences between the plankton and null distance distributions to assess whether planktonic organisms were closer or further apart than expected under a random distribution. The computed PNKS values were used as a measurement of association strength. These values were rescaled between 0 and 1, and made negative for cases of negative associations, i.e. when organisms were further away than expected.

#### Co-occurrence

Another association metric was computed based on the co-occurrence of organisms, computed is the correlation of their abundances across ISIIS images. By definition, this metric can only be computed across plankton groups. For each pair of plankton groups, the Pearson correlation of log-transformed concentrations was computed from 10,000 images. This process was repeated 100 times, and the results were averaged across repetitions. These numbers were shown to provide consistent results. Only significant correlations (p < 0.01) were retained.

#### Size-based association

Finally, it was desirable to compare these metrics to “true” associations, such as those from an empirical network Freilich et al. (2018). However, the low taxonomic resolution of the dataset prevented such an approach. Instead, a size-based method was used to infer predator-prey relationships. For each pair of plankton groups, one taxon was designated as the predator and the other as the prey. From each group, 10,000 organisms were sampled and ESD was used to compute 10,000 predator-to-prey size ratios (PPSR). The proportion of PPSRs falling within an optimal range was used as a metric of potential predation pressure. Five optimal PPSR ranges were explored (Table S3), around common values for zooplankton Hansen et al. (1994). The one yielding values most similar to the distance-based association was retained. Unlike the distance-based and co-occurrence approaches, this metric was not symmetrical, and was thus made symmetrical by taking the maximum value for each pair. To distinguish between significant and non-significant associations, 10,000 PPSR values were computed by randomly sampling the dataset, establishing a probability threshold below which predation is less likely to occur than by chance.

#### Comparison of association metrics

The three different association metrics (distance-based, co-occurrence and size-based) were compared to assess whether they capture the same ecological information. For all metrics, we retained only significant associations. Values were rescaled to a range of -1 to 1, with positive values indicating positive associations and negative values indicating negative associations. A Mantel test, specifically designed for association matrices, was performed to compare these metrics. However, this test requires complete matrices with no missing values, a requirement that our metrics did not meet, particularly for the distance-based one where we excluded cases where fewer than 10,000 distances could be computed. To address this limitation, we performed two versions of the Mantel test: first by replacing all missing association values with zeros, and second by analyzing only the subset of plankton groups for which all three matrices were complete. As a complementary approach, we utilized the size-based metric as an a priori binary classifier, defining expected associations between plankton groups. We then evaluated how effectively the distance-based and co-occurrence metrics detected these expected associations by calculating precision and recall scores.

### Individual-based model

To interpret the observed distances between planktonic organisms, we developed a simple individual-based model simulating attraction between organisms. In this model, organisms are represented as points with 3D positions randomly drawn within the imaged volume (similar to our null dataset). Distances in 2D between these points are thus equivalent to the null distances presented above. We implemented a density-based attraction mechanism where each point moves toward areas of higher organism density. All computations were performed in 3D, though distances were measured in 2D to match our plankton image analysis. We simulated 1,000 images with a single displacement step per image. Motility rate (i.e. proportion of moving points) was set to 96% to match the expected motility in the plankton dataset based on taxon-specific displacement capabilities. Two model parameters were tuned through a grid-search procedure: the sensory radius (i.e. kernel for density computation) and swimming ability (i.e. displacement length). We tested three sensory radius values around the identified optimal distance threshold (5, 10, and 20 cm) and four swimming ability values (displacement of 1, 2, 4 or 8 mm), corresponding to 2-3 body lengths of organisms in the dataset. Parameter selection was based on minimizing the Kuiper statistic between 10,000 quantiles of post-displacement distances and observed plankton distances, calculated using 100 subsamples of 1,000 plankton images. Selected parameters were 10 cm for sensory radius and 2 mm for displacement length (Figure S5).

Finally, distances generated by the individual-based model were used to evaluate the potential impact of lower recall in the image analysis (i.e. failing to detect all organisms in a given group). To simulate varying recall levels, distances were repeatedly subsampled to reflect recall values ranging from 1 to 0.1 (recalls measured on independent test sets were ∼0.9 for the segmentation and ranged 0.4 to 0.8 depending on the group for the classification, Table S1). The results showed that PNKS was largely insensitive to changes in recall (Figure S9).

## Data and code availability

Code and raw data are made available online (https://doi.org/10.5281/zenodo.16903101) Panaïotis (2025).

## Acknowledgement

The authors thank the officers and crew of the R/V Tethys II, as well as the additional scientists who participated in the VISUFRONT cruise: R. K. Cowen, R. Faillettaz, C. M. Guigand, M. Lilley, F. Lombard and J. Luo. They also appreciate L. Caray-Counil, S. Gasparini and D. Altukhov for their assistance with the initial stages of image identification, as well as B. Woodward for his help in setting up the segmentation and classification algorithms. Special thanks to D. Mayor and K. Cook for their valuable suggestions, and to L. Guidi for the early intuition that ultimately led to this work. Data acquisition during the VISUFRONT cruise was funded by the Partner University Fund and supported by the French Oceanographic Fleet through ship time. This work was granted access to the HPC resources of IDRIS under the allocation 2021-AD011013092 made by GENCI. We are also grateful to the Roscoff Bioinformatics platform ABiMS (http://abims.sb-roscoff.fr), part of the Institut Français de Bioinformatique (ANR-11-INBS-0013) and BioGenouest network, for providing computing resources. We thank the EMBRC platform PIQv for image analysis: this work was supported by EMBRC-France, whose French state funds are managed by the ANR within the Investments of the Future program under reference ANR-10-INBS-02. This work was supported by projects WWWPIC funded by the Belmont Forum through the Agence Nationale de la Recherche ANR-18-BELM-0003-01; CALIPSO funded by Schmidt Sciences and BIOcean5D funded by Horizon Europe (101059915). Views and opinions expressed are those of the author(s) only and do not necessarily reflect those of the European Union. Neither the European Union nor the granting authority can be held responsible for them.

