## Supplementary material for "A closer look at plankton: potential interactions inferred from centimeter-scale in situ observations"

Thelma Panaïotis[1,2], Jean-Olivier Irisson[2], Mara Freilich[3], BB Cael[4]

1. National Oceanography Centre, SO143ZH Southampton, UK
2. Laboratoire d'Océanographie de Villefranche, Sorbonne Université, 06230 Villefranche-sur-Mer, France
3. Department of Earth, Environmental and Planetary Science, Division of Applied Mathematics, Brown University, 02912 Providence, RI, USA
4. Department of the Geophysical Sciences, University of Chicago, 60637 Chicago, IL, USA

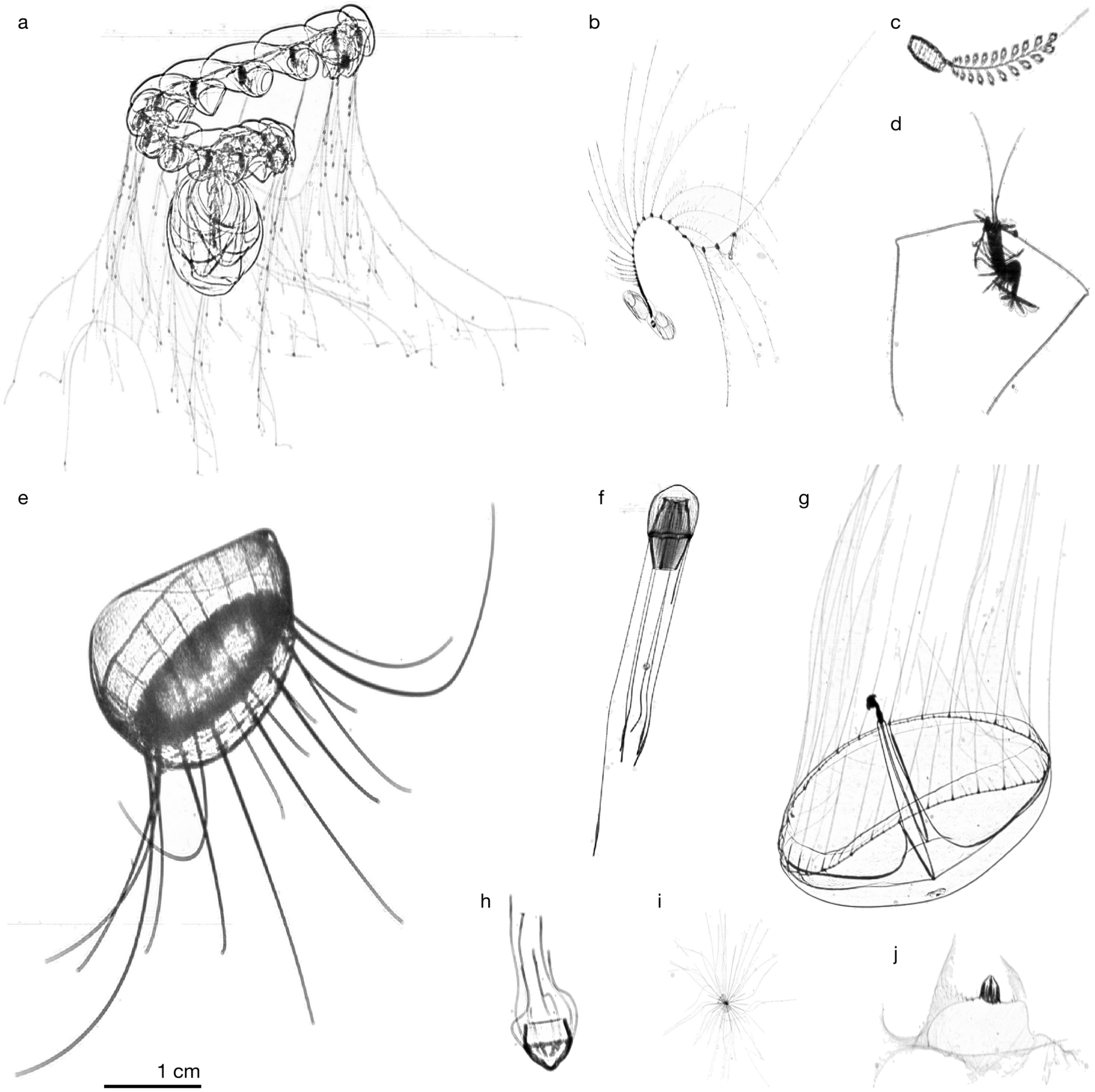


Figure S1: Examples of undisturbed delicate planktonic organisms imaged by ISIIS. (a) Prayidae (Siphonophorae), (b) Diphyidae (Siphonophorae), (c) Doliolida, (d) Eumalacostraca, (e) *Solmissus* (Cnidaria), (f) Scyphozoa (Cnidaria), (g) *Geryonia proboscidalis* (Cnidaria), (h) Scyphozoa (Cnidaria), (i) Foraminifera, (j) Cyddipida (Ctenophora).

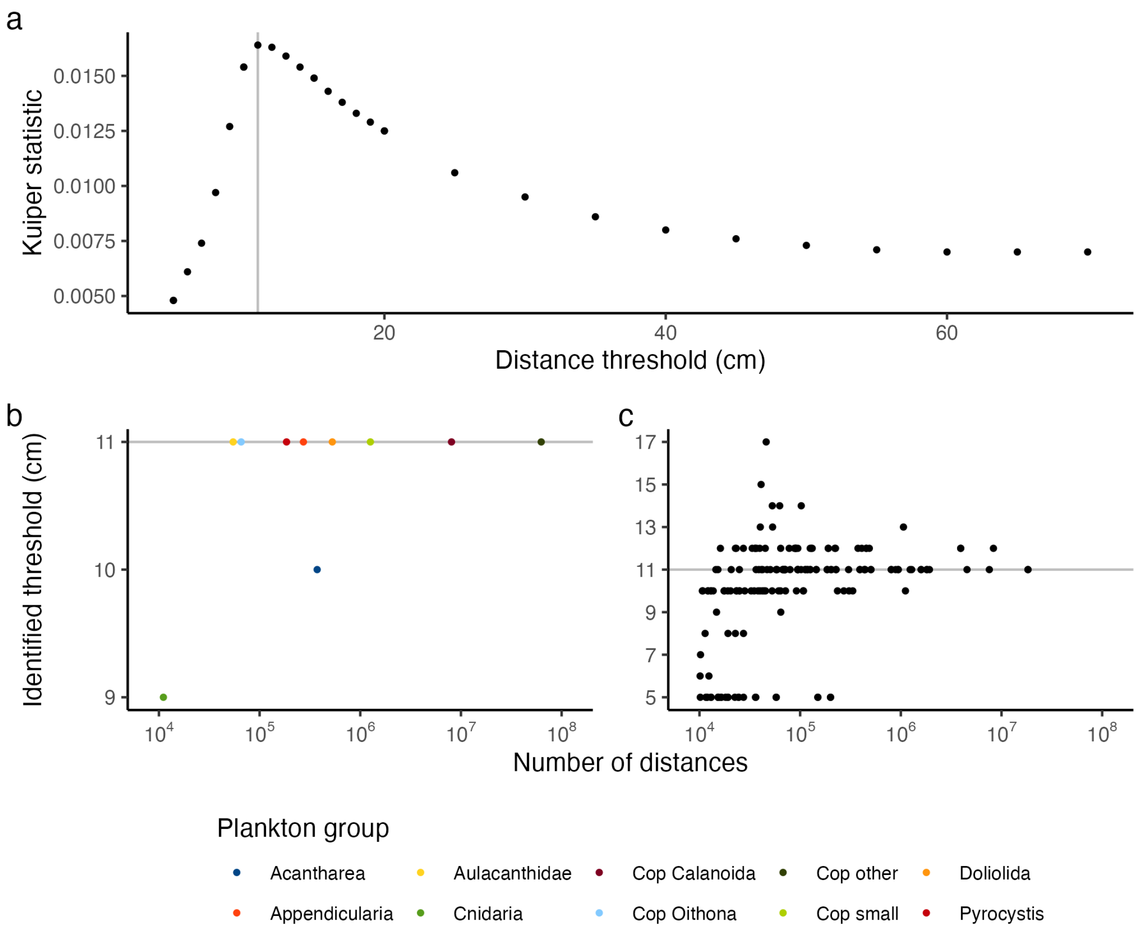


Figure S2: Identification of optimal distance ranges in terms of information content. (a) Plankton to null Kuiper statistic (PNKS) value for various distance thresholds, considering all planktonic organisms regardless of taxonomy. The vertical line highlights the highest PNKS value, indicating the most informative range of distances. (b, c) Identified thresholds versus number of retained distances for intra-group (b) and inter-group (c) distances. The gray horizontal line indicates the threshold selected for all analyses.

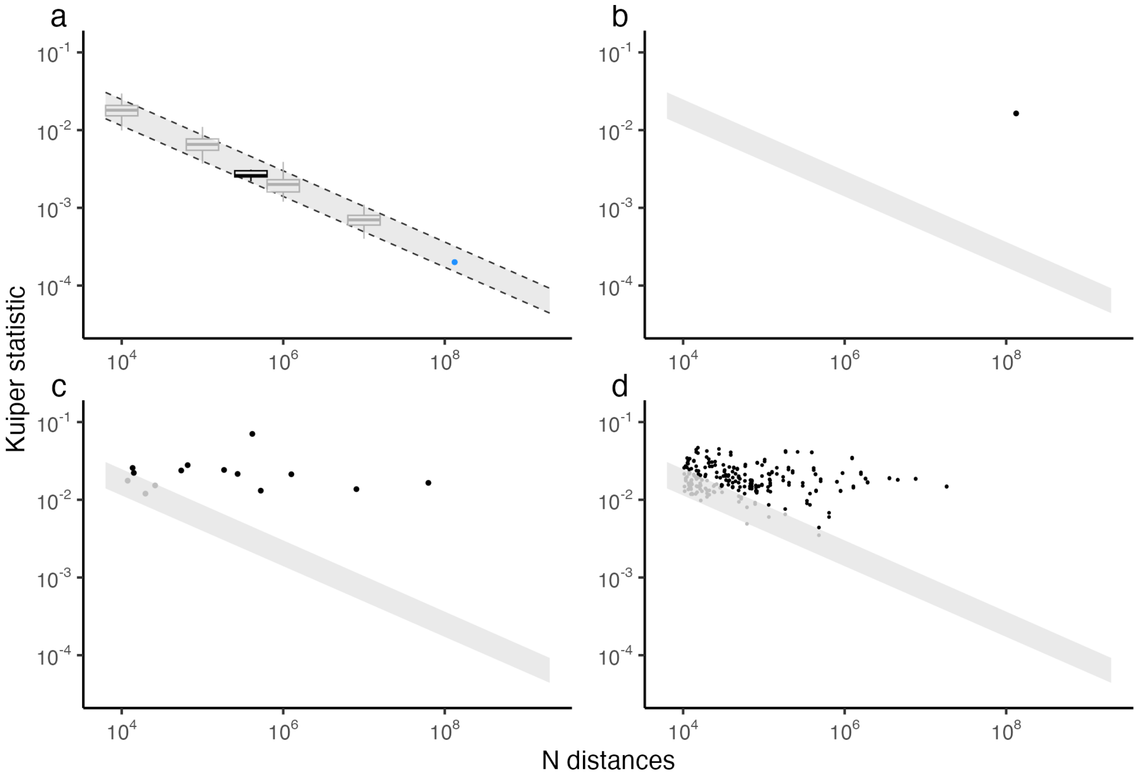


Figure S3: Kuiper statistics versus number of distances. (a) Null to null Kuiper statistics (NNKS) are shown as gray boxplots for subsamples of various sizes. Dashed lines represent the extrapolation of NNKS values for larger numbers of distances using quantile regression ($\tau$ = 0.05 and $\tau$ = 0.95), with the gray ribbon indicating the range of expected Kuiper statistic values under the null hypothesis (also reported in panels b–d), i.e. a distribution of distances not different from those expected under a random distribution. The black boxplot shows NNKS values from null datasets representing Acantharea only. Blue dots (overlaid) show NNKS values from three additional null datasets with a larger number of distances. (b) Plankton to null Kuiper statistic (PNKS) for all plankton distances. (c) PNKS for intra-group distances. (d) PNKS for inter-group distances.

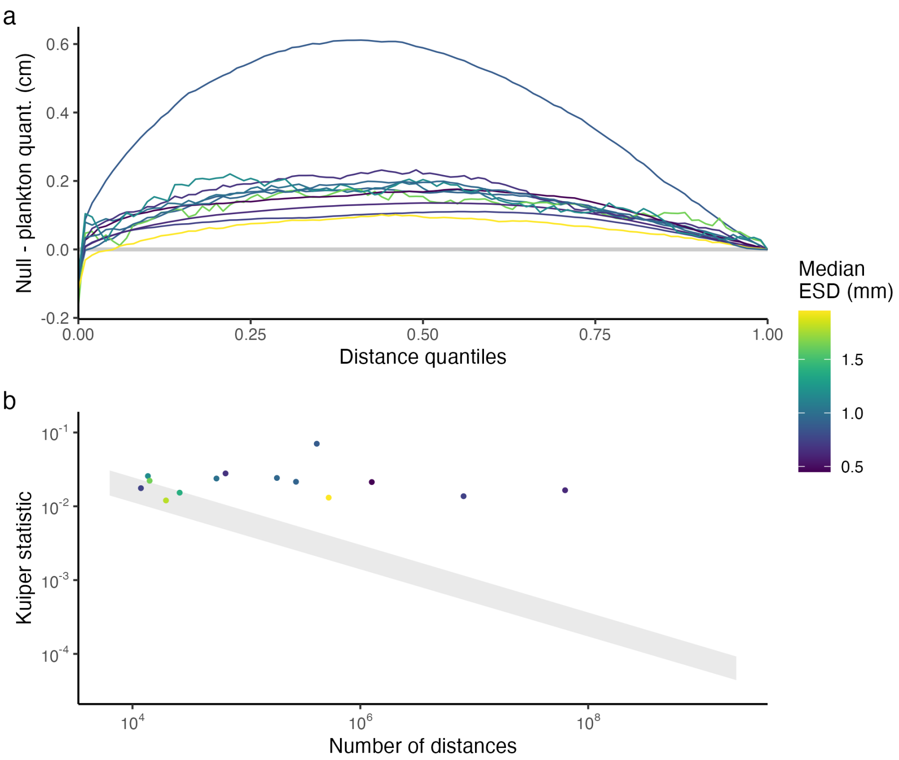


Figure S4: Absence of relationships between organisms size and observed distances. (a) Difference between quantiles of expected distances (random distribution) and quantiles of observed plankton distances, plotted against distance percentiles, colored by median equivalent spherical diameter (ESD) for each plankton group. (b) PNKS values for intra-group distances, colored by ESD for each plankton group.

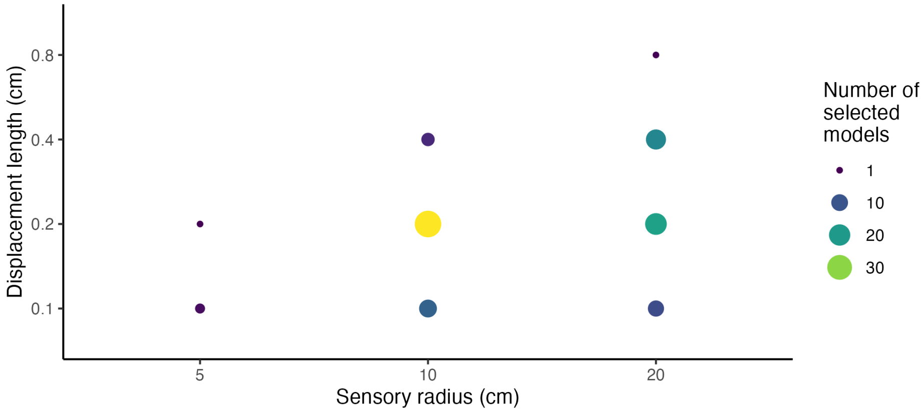


Figure S5: Number of times each set of hyperparameters was selected across 100 agent-based model runs. For each run, 12 parameter sets (3 sensory radius values and 4 displacement length values) were tested, and the set that best reproduced the observations was selected. For the final model, a sensory radius of 10 cm and a displacement length of 2 mm were selected.

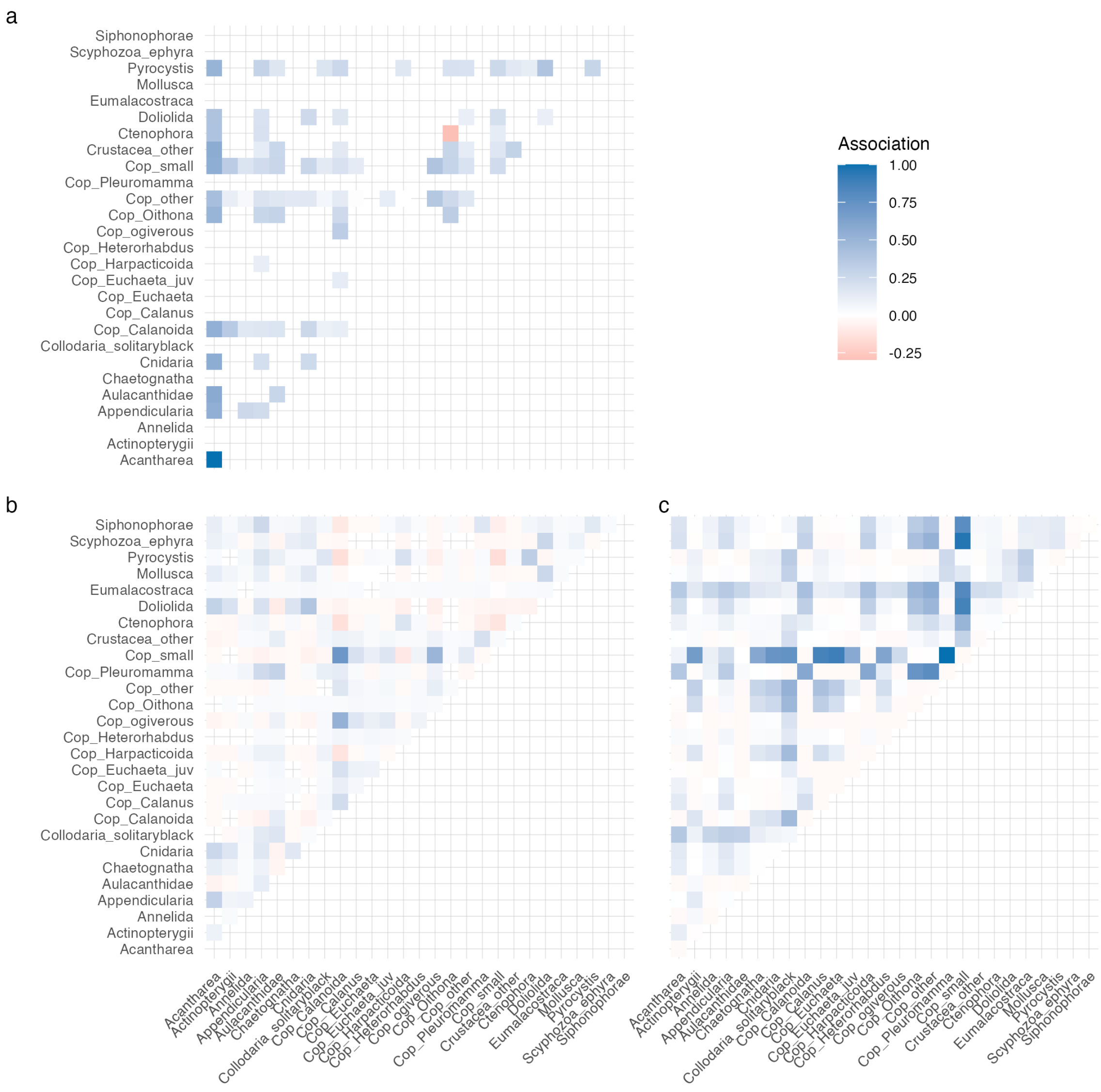


Figure S6: Association matrices for (a) the distance-based metric, (b) the co-occurrence metric and (c) the size-based metric.

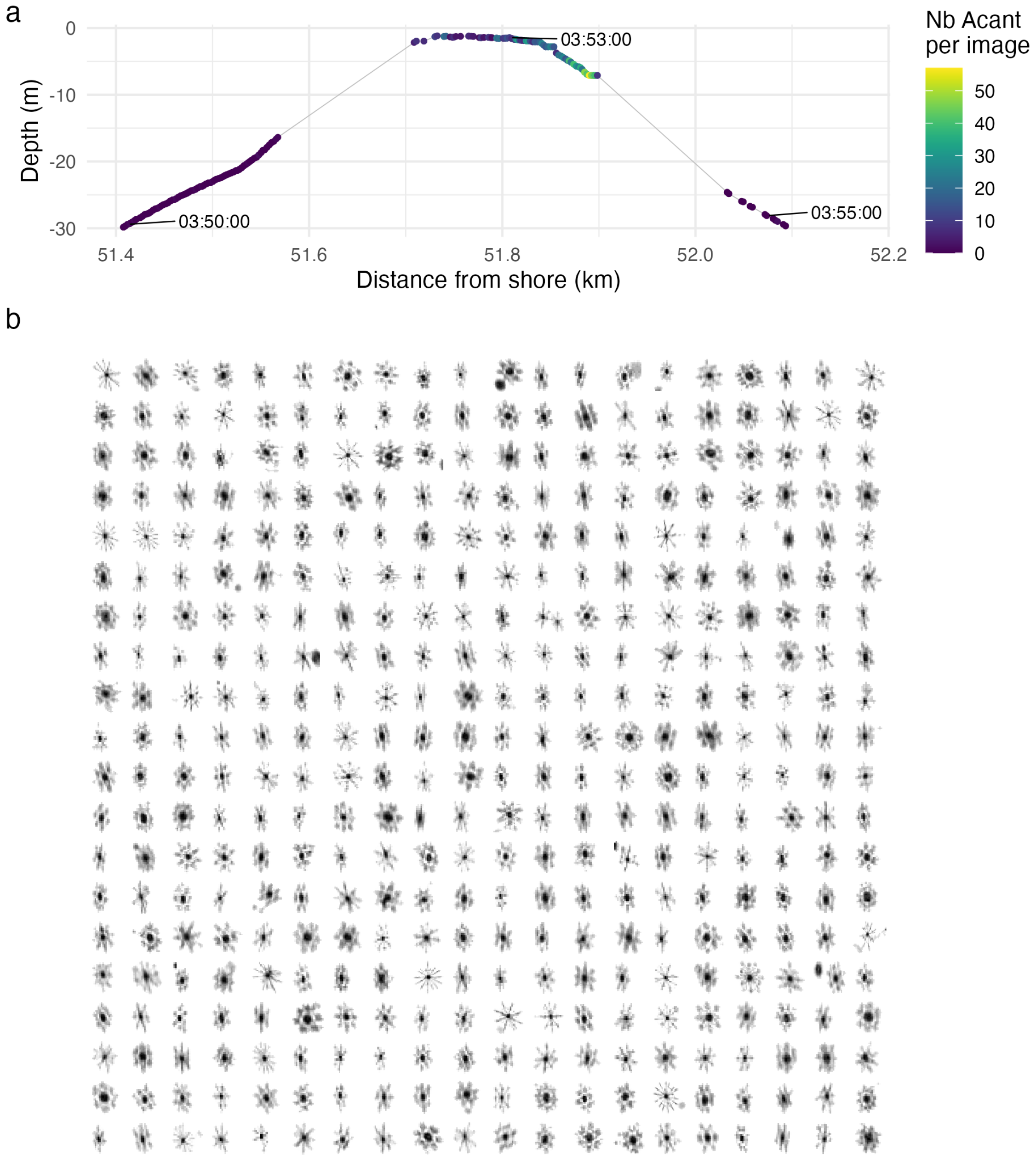


Figure S7: (a) Illustration of ISIIS sampling (continuous gray line) crossing a patch of Acantharea between the surface and 10 meter depth. Each point corresponds to an image (~50 cm long) and is coloured according to the number of Acantharea, showing a sharp increase within the patch. This sequence corresponds to ~5 minutes of sampling. (b) Mosaic of 400 randomly selected objects automatically classified as Acantharea from within such patches. Nearly all were true Acantharea, ruling out misclassification artifacts as an explanation for observed distances.

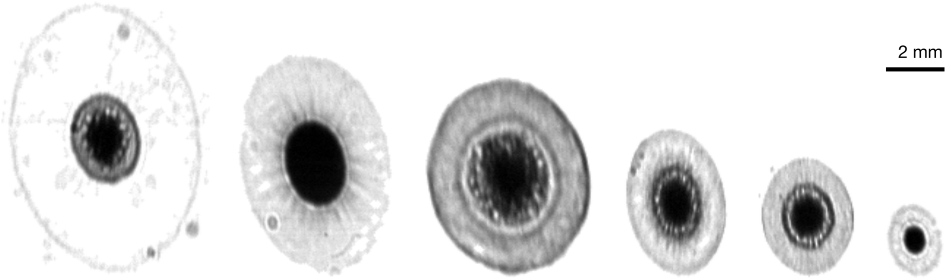


Figure S8: Examples of distorted spherical solitary Collodaria of various sizes resulting from variations in towing speed. A correction factor was calculated as the median ratio of height to width and applied globally to adjust the horizontal positions of all planktonic organisms.

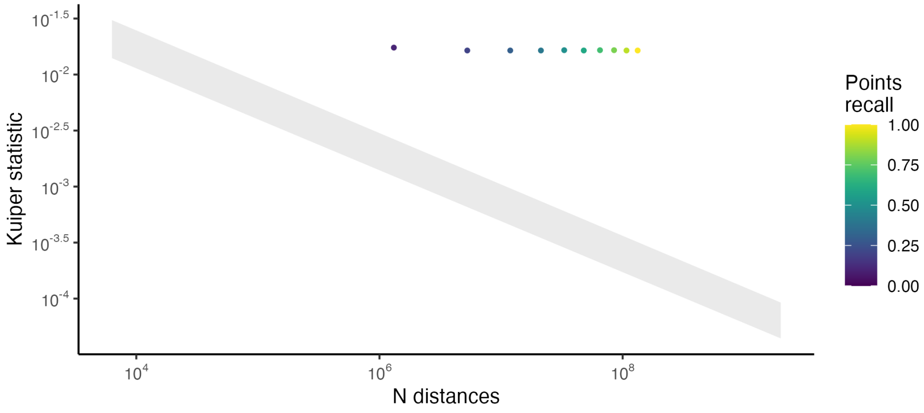


Figure S9: Effect of lower recall values on Kuiper statistic. Using distances obtained from the agent-based model simulating plankton distances, distances were subsampled to mimic lower recall values (i.e. non detected planktonic organisms). The grey ribbon indicates the range of expected Kuiper statistic values under the null hypothesis, as explained in Figure S3.

Table S1: Plankton groups included in this study, as well as classification precision and recall assessed on an independent test set, and the number of objects in each group. Copepoda was initially sorted as a unique group but was later partially further refined into 11 subgroups.

| Plankton group | Precision | Recall | N objects |
|---|---|---|---|
| Acantharea | 0.96 | 0.43 | 270,117 |
| Actinopterygii | 0.81 | 0.45 | 21,652 |
| Annelida | 0.83 | 0.68 | 57,660 |
| Appendicularia | 0.89 | 0.46 | 739,224 |
| Aulacanthidae | 0.98 | 0.64 | 272,589 |
| Chaetognatha | 0.62 | 0.62 | 23,592 |
| Cnidaria | 0.79 | 0.73 | 41,609 |
| Collodaria_solitaryblack | 0.93 | 0.76 | 64,168 |
| Copepoda | 0.99 | 0.52 | |
| Cop_Calanoida | - | - | 2,441,290 |
| Cop_Calanus | - | - | 41,104 |
| Cop_Euchaeta | - | - | 15,510 |
| Cop_Euchaeta_juv | - | - | 23,697 |
| Cop_Harpacticoida | - | - | 139,048 |
| Cop_Heterorhabdus | - | - | 7,707 |
| Cop_ogiverous | - | - | 19,653 |
| Cop_Oithona | - | - | 274,844 |
| Cop_other | - | - | 10,791,307 |
| Cop_Pleuromamma | - | - | 506 |
| Cop_small | - | - | 1,233,356 |
| Crustacea_other | 0.69 | 0.53 | 190,665 |
| Ctenophora | 0.71 | 0.70 | 155,447 |
| Doliolida | 0.89 | 0.80 | 155,674 |
| Eumalacostraca | 1.00 | 0.40 | 6,781 |
| Mollusca | 0.63 | 0.56 | 23,830 |
| Pyrocystis | 0.94 | 0.63 | 421,922 |
| Scyphozoa_ephyra | 1.00 | 0.44 | 13,568 |
| Siphonophorae | 0.87 | 0.76 | 133,756 |

Table S2: Mantel test results on distance matrices obtained using three different association matrices. The Mantel statistic was based on Spearman's rank correlation. For the zero-filled method, any pair of plankton groups for which a metric could not be computed was assigned a value of zero. For the reduced methods, plankton groups for which a given metric could not be computed were successively removed from the matrices to achieve complete matrices. All Mantel statistics were found to be non-significant.

| Compared matrices | Mantel statistic | |
|---|---|---|
| | Zero-filled | Reduced |
| Size-based VS co-occurrence | -0.04 | -0.25 |
| Size-based VS distance-based | -0.05 | -0.10 |
| Co-occurrence VS distance-based | 0.01 | -0.30 |

Table S3: Explored predator to prey size ratios (PPSR) ranges, selected to span five center values between 2 and 10 on a log-scale.

| **Range** | **Centre** |
|---:|---:|
| [5, 20] | 10 |
| [4, 16] | 8 |
| [3, 12] | 6 |
| [4, 8] | 4 |
| [1, 4] | 2 |
